# High residual prevalence of vaccine-serotype *Streptococcus pneumoniae* carriage after introduction of a pneumococcal conjugate vaccine in Malawi: a prospective serial cross-sectional study

**DOI:** 10.1101/445999

**Authors:** Todd D. Swarthout, Claudio Fronterre, José Lourenço, Uri Obolski, Andrea Gori, Naor Bar-Zeev, Dean Everett, Arox W. Kamng’ona, Thandie S. Mwalukomo, Andrew A. Mataya, Charles Mwansambo, Marjory Banda, Sunetra Gupta, Peter Diggle, Neil French, Robert S. Heyderman

## Abstract

**Background:** There are concerns that pneumococcal conjugate vaccines (PCV) in sub-Saharan Africa sub-optimally interrupt vaccine-serotype (VT) carriage and transmission, thus limiting vaccine-induced direct and indirect protection. We assessed carriage in vaccinated children and unvaccinated populations targeted for indirect protection, between 4 and 7 years after Malawi’s November 2011 introduction of PCV13 using a 3+0 schedule.

**Methods:** We conducted sequential prospective nasopharyngeal carriage surveys between 2015 and 2018 among healthy PCV-vaccinated and PCV-unvaccinated children, and HIV-infected adults. VT and NVT carriage risk by age was analysed by non-linear regression.

**Results:** Among PCV-vaccinated children, there was a 24% relative reduction in carriage, from a mean 21.1% to 16.1%; 45% reduction among older PCV-unvaccinated children, from 27.5% to 15.2%; 41.4% reduction among adults, from 15.2% to 8.9%. Using carriage data from children 3.6 to 10 years of age, VT carriage probability declined with age, with a similar prevalence half-life among PCV-vaccinated (3.34 years) and PCV-unvaccinated (3.26 years) children.

**Conclusion:** Compared to high-income settings, the 3+0 schedule in Malawi has led to a sub-optimal reduction in pneumococcal carriage prevalence. This is likely due to recolonisation of vaccinated children with waning vaccine-induced immunity, resulting in insufficient indirect protection of unvaccinated populations. Rigorous evaluation of strategies to augment vaccine-induced control of carriage, including alternative schedules and catch-up campaigns is required.

## BACKGROUND

*Streptococcus pneumoniae* is estimated to be responsible for over 500 000 deaths every year in children aged 1 to 59 months worldwide, with the highest burden among African children.^1^ *S. pneumoniae* has over 90 immunological serotypes and is a common coloniser of the human nasopharynx, particularly in young children, resource-poor and HIV-affected populations.^1^ Although most carriers are asymptomatic, pneumococcal colonisation is a necessary prerequisite for transmission and the development of pneumonia, meningitis, and bacteraemia.^2^

In Europe and North America, routine infant administration of pneumococcal conjugate vaccine (PCV) has rapidly reduced vaccine-serotype (VT) invasive pneumococcal disease (IPD) and carriage.^3–6^ Importantly, this has occurred in vaccinated and unvaccinated age groups. Thus, indirect protection resulting from a reduction in carriage and transmission amplifies PCV impact and cost-effectiveness.^7^ Pneumococcal epidemiology in sub-Saharan Africa is characterised by high rates of carriage and transmission, differing markedly from high-income settings.^8,9^ Carriage studies pre-dating PCV introduction in Kenya,^8^ Mozambique,^10^ Malawi,^11^ The Gambia,^12^ and South Africa^13^, for example, reported VT carriage prevalences ranging from 49.7% to 28.2% in under 5s, with colonisation occurring rapidly early in life.^14^

Vaccine trials and post-routine-introduction studies in Africa have demonstrated substantial direct effects of PCV against IPD, pneumonia, and all-cause mortality among young children.^15–18^ Although Kenya,^19^ The Gambia,^18^ Mozambique^20^, and South Africa^21^ have reported VT carriage reductions, residual carriage prevalences are still higher than in industrialised countries.^22–24^ In addition, there is evidence of rapid onset of NVT replacement in the region.^25^ Thus, it is uncertain whether PCV introduction in sub-Saharan Africa will achieve the sustained control of pneumococcal carriage necessary to effectively interrupt transmission and so disease. This is of particular concern in many sub-Saharan African countries where the 3+0 schedule has been implemented into infant expanded programmes on immunisation (EPI).^26^

In November 2011, Malawi (previously PCV-naïve) introduced 13-valent PCV as part of the national EPI using a 3+0 schedule (6, 10 and 14 weeks of age). A three-dose catch-up vaccination campaign included infants <1 year of age. Field studies among age-eligible children have reported a high PCV13 uptake of 90–95%,^27,28^ even higher than the 83% previously reported by WHO/UNICEF.^29^ In 2011, Malawi adopted Option B+, whereby all HIV-positive pregnant or breastfeeding women commence lifelong full ART regardless of clinical or immunological stage, dramatically reducing mother-to-child-transmission.^30^

We hypothesised that despite evidence of PCV13 impact on IPD and pneumonia in Malawi,^31,32^ there would be persistent VT carriage and that this would maintain transmission in both childhood and adult reservoirs. We have investigated this among PCV13-vaccinated children (in whom vaccine-induced immunity wanes after the first year of life^33^); children too old to have received PCV13; and HIV-infected adults on antiretroviral therapy (ART) who do not routinely receive pneumococcal vaccination (previously demonstrated to have a high carriage prevalence^34,35^).

## METHODS

### Study Design

This was a prospective cross-sectional observational study using stratified random sampling to measure pneumococcal nasopharyngeal carriage in Blantyre, Malawi. Sampling consisted of a time series profile from twice-annual surveys over 3.5 years.

### Study population and recruitment

Blantyre is located in Southern Malawi with an urban population of approximately 1·3 million. Recruitment included four groups: i) healthy infants 4-8 weeks old prior to first dose of PCV, recruited from vaccination centres using systematic sampling; ii) randomly sampled healthy children 18 weeks–7 years old who received PCV as part of EPI or the catch-up campaign, recruited from households and public schools; iii) randomly sampled healthy children 3–10 years old who were age-ineligible (born on or before 11 November 2010 and therefore too old) to receive PCV as part of EPI or the catch-up campaign), recruited from households and public schools; and iv) HIV-infected adults 18–40 years old and on ART, recruited from Blantyre’s Queen Elizabeth Central Hospital ART Clinic using systematic sampling. Recruitment of infants 4-8 weeks was implemented starting survey-5. Recruitment of children 18 weeks - 2 years old was implemented starting survey-4, after evidence of persistent carriage among children 3-10 years older during the first three surveys. Exclusion criteria for all participants included current TB treatment, pneumonia hospitalisation ≤14 days before study enrolment or terminal illness. Exclusion criteria for children included reported immunocompromising illness (including HIV), having received antibiotics ≤14 days prior to screening, having received PCV if age-ineligible or not having received PCV if age-eligible. Individuals were not purposely resampled but were eligible if randomly re-selected in subsequent surveys.

### Site selection

Households, schools and vaccination centres were selected from within three non-administrative zones representative of urban Blantyre’s socioeconomic spectrum in medium- to high-density townships. These zones were further divided into clusters, allowing for approximately 25 000 adults per zone and 1 200 adults per cluster. Clusters were not purposely resampled but eligible if randomly selected in subsequent surveys. Within each cluster, after randomly choosing a first house, teams moved systematically, recruiting one eligible child per household until the required number of children were recruited from each cluster. Individual school-goers were randomly selected from school registers, and letters sent home inviting parents or legal guardians to travel to the school to discuss the study and consider consenting to their child’s participation.

### Determining PCV vaccination status

A child was considered “PCV-vaccinated” if s/he had received at least one dose of PCV prior to screening. Vaccination status and inclusion/exclusion criteria were further assessed from subject-held medical records (known as Health Passports). If a child was reported by the parent/guardian to be PCV-vaccinated but no health passport was available, a questionnaire was applied. The questionnaire was developed by identifying, among a subset of 60 participants, four questions most commonly answered correctly by parents/guardians of children with proof of PCV vaccination. The questions included child’s age when vaccinated, vaccine type (oral or injectable), anatomical site of vaccination, and which other (if any) vaccines were received at the time of PCV vaccination. If the child was PCV age-eligible and answered all four questions correctly, the child was recruited as “PCV-vaccinated.”

### Sample size

The sample size strategy was a pragmatic approach to allow for adequate precision of the carriage prevalence estimates. Using VT carriage as the primary endpoint, the sample size was calculated based on the precision of the prevalence estimation, assuming an infinite sampling population. Among children 3–7 years old (vaccinated), an absolute VT prevalence up to 10% was expected, with a sample of 300/survey providing a 95% confidence interval (CI) of 6·6–13·4%. Among children 3–10 years old (unvaccinated) and HIV-infected adults, an absolute VT prevalence of 20% was expected, with a sample of 200/survey providing a 95% CI of 14·5–25·5%.

### Nasopharyngeal swab collection

A nasopharyngeal swab (NPS) was collected from each participant using a nylon flocked swab (FLOQSwabs™, Copan Diagnostics, Murrieta, CA, USA) and then placed into 1·5mL skim milk-tryptone-glucose-glycerol (STGG) medium and processed at the Malawi–Liverpool–Wellcome Trust (MLW) laboratory in Blantyre, according to WHO recommendations.^36^ Samples were frozen on the same day at −80°C.

### Pneumococcal identification and latex serotyping

After being thawed and vortexed, 30 μL NPS–STGG was plated on gentamicin-sheep blood agar (SBG; 7% sheep blood agar, 5 μl gentamicin/mL) and incubated overnight at 37°C in 5% CO2. Plates showing no *S. pneumoniae* growth were incubated overnight a second time before being reported as negative. *S. pneumoniae* was identified by colony morphology and optochin disc (Oxoid, Basingstoke, UK) susceptibility. The bile solubility test was used on isolates with no or intermediate (zone diameter <14mm) optochin susceptibility. A single colony of confirmed pneumococcus was selected and grown on a new SBG plate as before. Growth from this second plate was used for serotyping by latex agglutination (ImmuLex™ 7-10-13-valent Pneumotest; Statens Serum Institute, Denmark). This kit allows for differential identification of each PCV13 VT but not for differential identification of NVT serotypes; NVT and non-typeable isolates were therefore reported as NVT. Samples were batch tested on a weekly basis, blinded to the sample source. Latex serotyping results showed good concordance with whole genome sequence and DNA microarray serotyping.^37^

### Statistical analysis

Participant demographic characteristics were summarised using means, standard deviations, medians, and ranges for continuous variables and frequency distributions for categorical variables. Non-ordinal categorical variables were assessed as indicators. Carriage prevalence ratios (PR) were calculated over the study duration by log-binomial regression using months (30.4 days) between study start and participant recruitment, coded as a single time variable, allowing an estimate of prevalence ratio per month. Potential confounders were identified by testing the association between variables and included in the multivariable models when p<0.1. Adjusted prevalence ratios (aPR) were calculated using log-binomial regression. Confidence intervals are binomial exact. Statistical significance was inferred from two-sided p<0·05. Statistical analyses were completed using Stata 13.1 (StataCorp, College Station, TX, USA).

### Non-linear regression analysis for VT carriage decay rate and half-life

To better understand the rate at which VT and NVT carriage prevalence was decreasing, we developed a non-linear model to describe the variation in probability of VT or NVT carriage with age, adjusted for age at recruitment. The model is fitted using carriage data from children 3.6 to 10 years of age, maximising overlap of empirical data and allowing direct comparison of parameters between vaccinated and unvaccinated children. Model outputs were transformed into a population-level (decay) half-life (i.e. time in years for carriage prevalence in the sampled cohort to reduce to one-half of its peak) was log(2)/δ), where δ = rate of decay of VT or NVT carriage prevalence with age. Model parameters were estimated by maximum likelihood, and 95% confidence bands for the predicted exponential decay curves are obtained through parametric bootstrap. This analysis used R open-source software (www.r-project.org). Details of the analysis framework are in Supplement 1.

## RESULTS

Between 19 June 2015 and 6 December 2018, seven cross-sectional surveys were completed: dates for each survey were, respectively, (1) June–August, 2015; (2) October, 2015–April, 2016; (3) May–October, 2016; (4) November, 2016–April, 2017; (5) May–October 2017; (6) November, 2017–June, 2018; (7) June–December, 2018. 7554 individuals were screened (Figure 1), including 371 PCV-unvaccinated infants 4-8 weeks old, 602 PCV-vaccinated children 18 weeks–1-year old, 538 PCV-vaccinated children 2 years old, 2696 PCV-vaccinated children 3–7 years old, 1505 PCV-unvaccinated children 3–10years old, and 1842 HIV-infected adults 18–40 years old and on ART (PCV-unvaccinated). Among these, 24 (6·5%) infants prior to PCV vaccination, 196 (5·1%) children age-eligible for PCV, 96 (6·4%) children age-ineligible for PCV, and 67 (3·6%) adults were excluded (figure 1) from recruitment after screening. Twenty-three (23) participants (eighteen children, five adults) did not allow a swab to be collected after recruitment. The final analysis included 7148 participants: 346 infants recruited prior to first dose PCV, 566 children 18 weeks–1-year-old and PCV-vaccinated, 499 children 2 years old and PCV-vaccinated, 2565 children 3–7 years and PCV-vaccinated, 1402 children 3–10 years old and PCV-unvaccinated, and 1770 HIV-infected adults on ART and PCV-unvaccinated. Among the children in the final analysis, 3605 were recruited from households and 1427 from schools.

**Figure 1.**
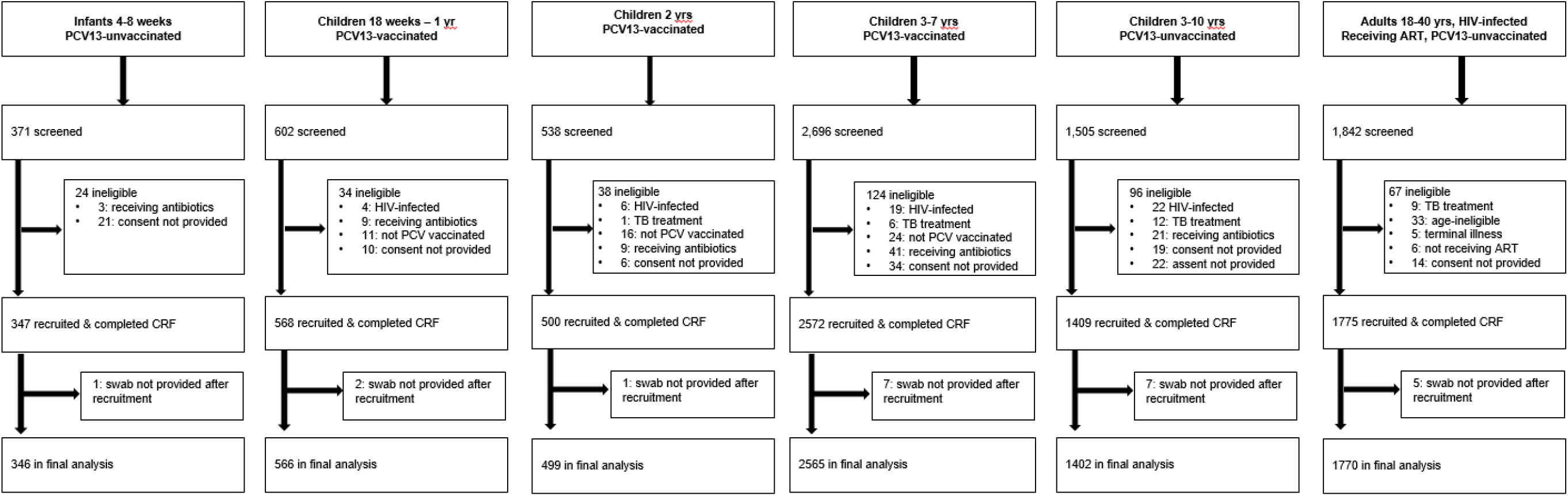
Screening, reasons for exclusion and recruitment

### Demographics and vaccination history

The surveyed groups had similar demographics (Table 1). However, a higher proportion of younger children 18 weeks–7-years-old (vaccinated) lived in houses with some lower infrastructure standards (walls and latrine facilities), relied more on shared communal water sources and scored lower on the aggregate index of household possessions. Among those screened and age-eligible for PCV vaccination, 98.7% (3785 / 3836) reported receiving at least one dose of PCV.

**Table 1:**
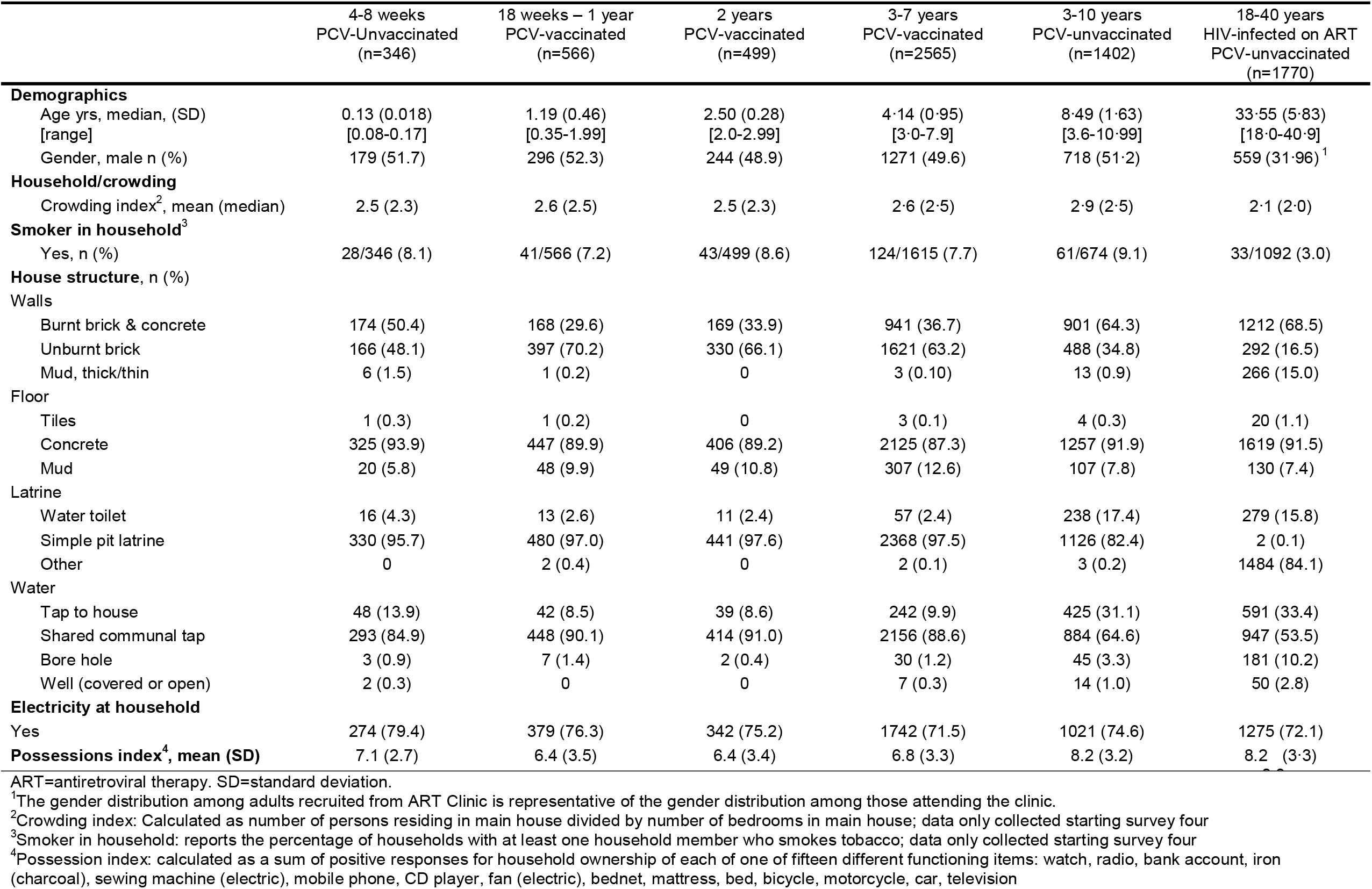
Demographic and household characteristics of child and adult participants

Among the 3630 PCV-vaccinated children recruited and providing an NPS, 1209 (33·3%) had documented (health passport) vaccination status with dates of vaccination; ranging from 86.5% among the youngest vaccinated age group (18 weeks–1-year-old) to 25.7% among the oldest vaccinated age group (3-7 years old). Among those with health passports confirming dates of vaccination, the median (IQR) age at first, second, and third dose of PCV were 6·3 (3·2), 11·2 (5·0), and 16·4 (8·1) weeks, respectively; 1143 (94·5%) received three doses PCV, 24 (2.0%) only two doses and 42 (3·5%) only one dose.

### Pneumococcal carriage

Among children 4-8 weeks (prior to first dose PCV) aggregated (surveys 5–7) VT and NVT carriage prevalence were respectively, 8·4% (95% CI 5·7–11.8) and 33·8% (95% CI 28·8–39·1). There was a 25·2% relative reduction in VT carriage, from 10 7% (95% CI 6·3–17·7) in survey 5 to 8·7% (4·4–16·5) in survey 7. (Figure 2; Table 2) When adjusted for age at recruitment, the adjusted prevalence ratio (aPR) over the 3 surveys was 0 974 (95% CI 0·902–1·052) p=0·506. Supplement 3 shows the aPR for VT carriage, stratified by individual survey within each age group. There was a 51·9% relative increase in NVT carriage, from 32·2% (24·5–41·1) in survey 5 to 48·9% (38·8–59·1) in survey 7. The aPR was 1·037 (1·002–1·073), p=0·038.

**Figure 2.**
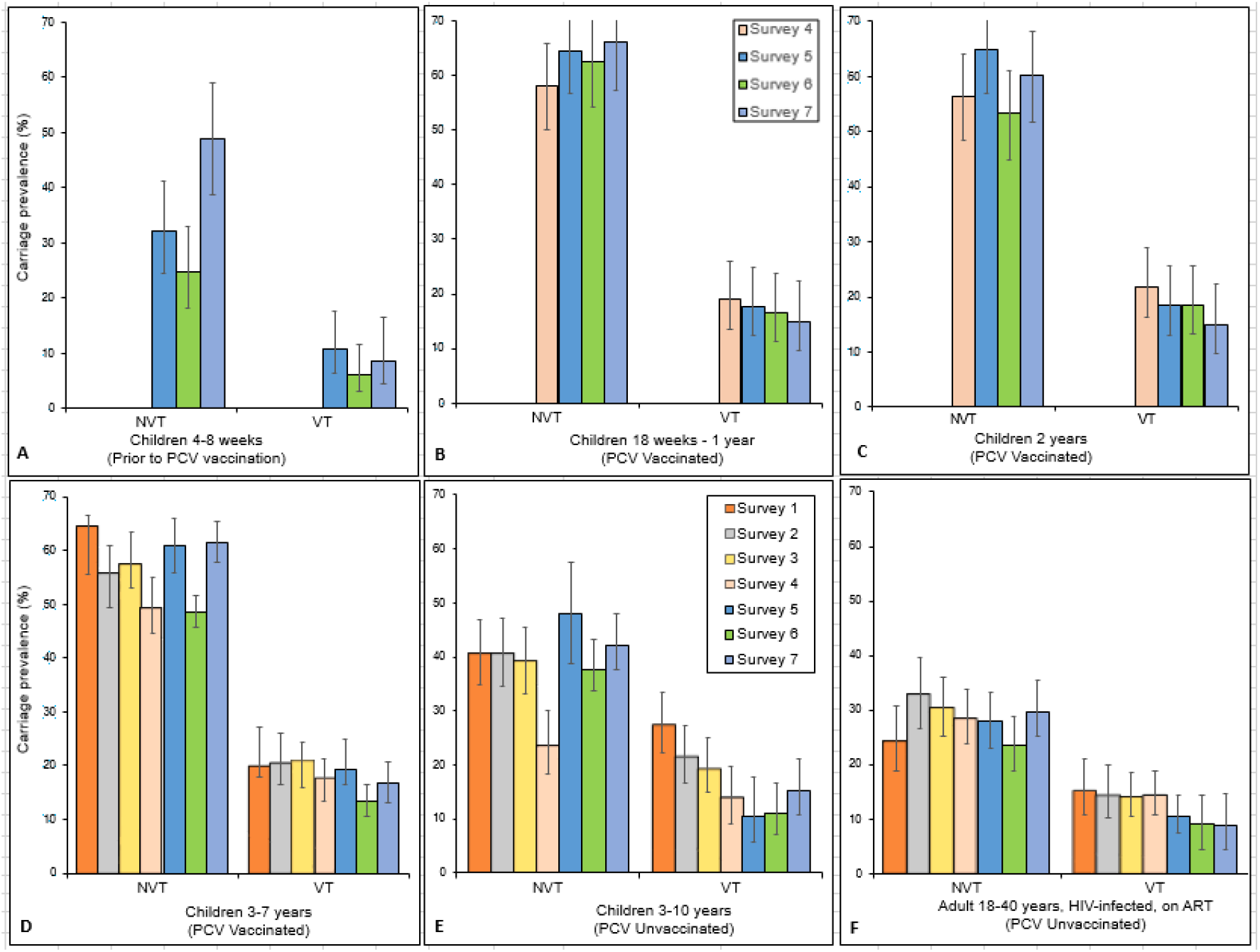
Prevalence of *Streptococcus pneumoniae* carriage per survey, stratified by study group and PCV vaccination status. Younger target groups were recruited starting survey 4 or 5, including children 4-8 weeks prior to their first dose PCV (A, surveys 5-7), PCV-vaccinated children 18 weeks–1-year old (B, surveys 4-7), and 2 years old (C, surveys 4-7). Older age groups, with data from surveys 1–7 included PCV-vaccinated children 3–7yrs (D), PCV-unvaccinated children 3–10yrs old (E), and HIV-infected adults on ART (F). 95% confidence interval error bars are shown. Prevalence of non-carriers is calculated by 1-(NVT+VT). Refer to Table 2 for VT & NVT prevalence stratified by survey and (adjusted) prevalence ratios; Refer to Appendix 2 for Proportion and frequency of VT carriage attributed to each VT, stratified by study.

**Table 2:**
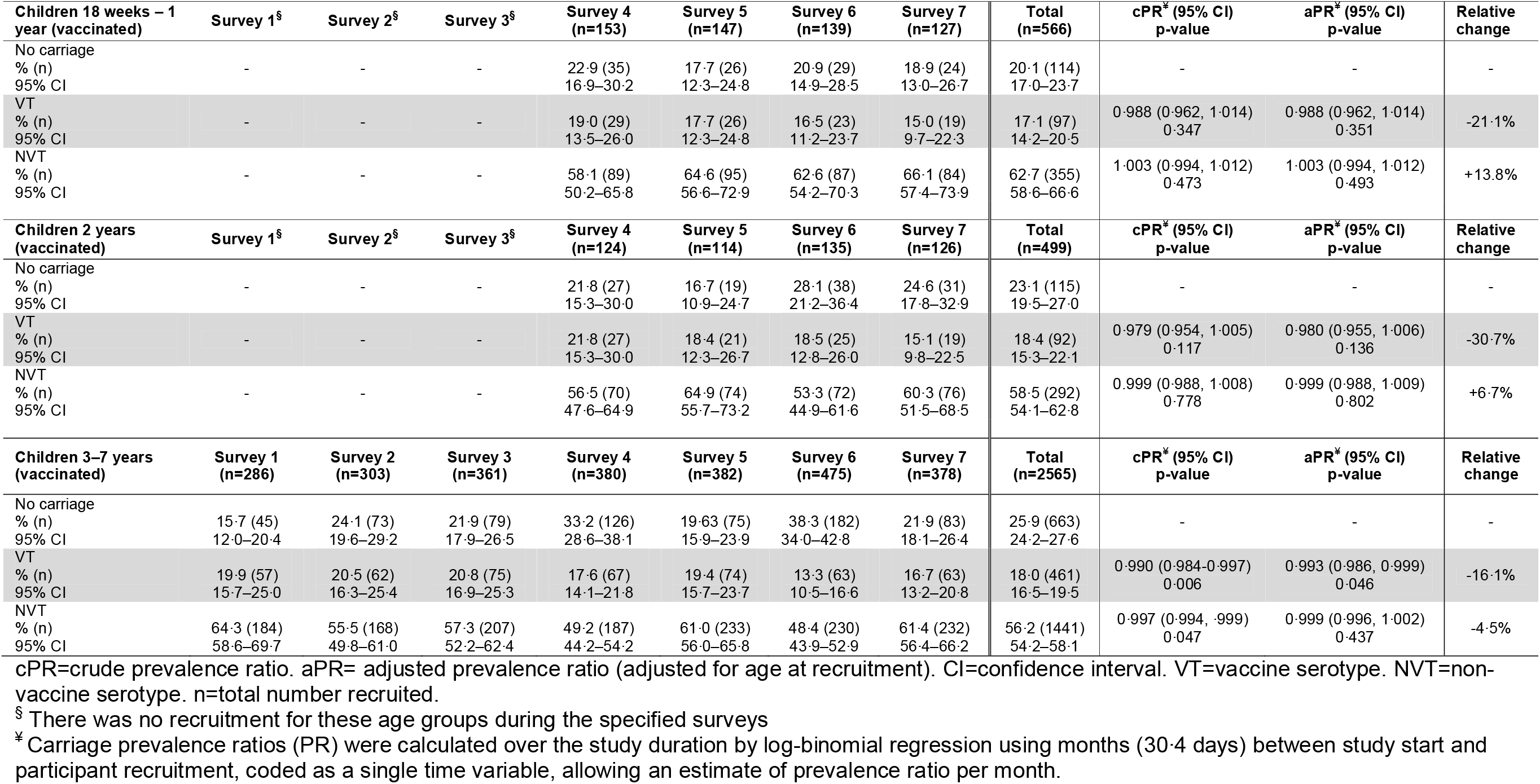
Vaccine and non-vaccine serotype *S. pneumoniae* carriage prevalence among PCV13-vaccinated study groups, stratified by survey, with prevalence ratios and relative change

Among PCV-vaccinated children 18 weeks–1-year-old (Figure 2; Table 2), aggregated (surveys 4–7) VT and NVT carriage prevalence were respectively, 17·1% (95% CI 14·2–20·5) and 62·7% (95% CI 58·6–66·6). There was a 21·1% relative reduction in VT carriage, from 19·0% (95% CI 13·5–26·0) to 15·0% (9·7–22·3); aPR 0·988 (95% CI 0·962–1·014), p=0·351. There was a 13·8% relative increase in NVT carriage, from 58·1% (50·2–65·8) to 66·1% (57·4–73·9); aPR: 1·003 (0·994–1·012), p=0·493.

Among children 2 years old (PCV-vaccinated), aggregated (surveys 4–7) VT and NVT carriage prevalence were respectively, 18·4% (95% CI 15·3–22·1) and 58·5% (95% CI 54·1–62·8). There was a 30·7% relative reduction in VT carriage, from 21·8% (15·3–30·0) to 15·1% (9·8–22·5); aPR: 0·980 (0·955–1·006), p=0·136. There was a 6·7% relative increase in NVT carriage, from 56·5% (47·6–64·9) to 60·3% (51·5–68·5); aPR: 0·999 (0·988–1·009), p=0·802. (Figure 2; Table 2)

Among children 3–7 years (PCV-vaccinated), aggregated (surveys 1–7) VT and NVT carriage prevalence were respectively, 18·0% (95% CI 16·5–19·5) and 56·2% (95% CI 54·2–58·1). There was a 16·1% relative reduction in VT carriage, from 19·9% (15·7–25·0) to 16·7% (13·2–20·8); aPR: 0·993 (0·986–0·999), p=0·046. (Figure 2; Table 2) A sensitivity analysis among the PCV-vaccinated age groups showed neither the overall VT prevalence nor the VT distribution changed significantly when limiting these analyses to children i) who received only one, only two, or all three doses PCV; ii) with document-confirmed PCV vaccination or iii) who adhered to the vaccination schedule to within 2 weeks of each scheduled dose (data not shown). There was a 4·5% relative reduction in NVT carriage, from 64·3% (95% CI 58·6–69·7) to 61·4% (95% CI 56·4–66·2); aPR: 0·999 (0·996–0·1.002), p=0·437.

When stratified by age (in years) and aggregating survey data, reduction in VT carriage was not exponential among vaccinated children (Supplement 1). Though not statistically significant, VT carriage increased slightly during the first 4 years of life, from 16·6% (95% CI 11·9–22·6) among children 18 weeks-11 months old to 17·4% (13·6–21·7), 18·6% (15·1–22·1), and 19·5% (17·3–22·1) among 1-, 2-, and 3-year old children, respectively. VT carriage then decreased to 18·5% (16·0–20·9), 14·4% (10·8–19·0), 12·0% (6·7–19·3), and 7·0% (1·4–19·1) among 4-, 5-, 6-, and 7-year olds, respectively.

Among PCV-unvaccinated children 3–10 years old, aggregated (surveys 1–7) VT and NVT carriage prevalence were respectively, 18·2% (95% CI 16·2–20·3) and 38·5% (95% CI 35·9–41·0). There was a 44·7% relative reduction in VT carriage, from 27·5% (22·3–33·3) to 15·2% (10·8–21·0); aPR: 0·989 (0·979–0·999), p=0·029. There was a 3·2% relative increase in NVT carriage, from 40·8% (34·9–46·9) to 42·1% (35·4–49·2); aPR: 1·005 (0·999–1·010), p=0·101. (Figure 2; Table 3)

**Table 3:** Vaccine and non-vaccine serotype *S. pneumoniae* carriage prevalence among PCV13-unvaccinated study groups, stratified by survey, with prevalence ratios and relative change

| Children 4-8 weeks (unvaccinated) | Survey 1<sup>§</sup> | Survey 2<sup>§</sup> | Survey 3<sup>§</sup> | Survey 4<sup>§</sup> | Survey 5 (n=121) | Survey 6 (n=133) | Survey 7 (n=92) | Total (n=346) | cPR* (95% CI) p-value | aPR* (95% CI) p-value | Relative change |
| --- | --- | --- | --- | --- | --- | --- | --- | --- | --- | --- | --- |
| No carriage |  |  |  |  |  |  |  |  |  |  |  |
| % (n) | - | - | - | - | 57.0 (69) | 69.2 (92) | 42.4 (39) | 57.8 (200) | - | - | - |
| 95% CI |  |  |  |  | 48.0–65.6 | 60.8–76.5 | 32.7–52.8 | 52.4–63.1 |  |  |  |
| VT |  |  |  |  |  |  |  |  |  |  |  |
| % (n) | - | - | - | - | 10.7 (13) | 6.0 (8) | 8.7 (8) | 8.4 (29) | 0.973 (0. 901, 1.052) | 0.974 (0. 902, 1.052) | -25.2% |
| 95% CI |  |  |  |  | 6.3–17.7 | 3.0–11.6 | 4.4–16.5 | 5.7–11.8 | 0.493 | 0.506 |  |
| NVT |  |  |  |  |  |  |  |  |  |  |  |
| % (n) | - | - | - | - | 32.2 (39) | 24.8 (33) | 48.9 (45) | 33.8 (117) | 1.035 (1.001, 1.072) | 1.037 (1.002, 1.073) | +51.9% |
| 95% CI |  |  |  |  | 24.5–41.1 | 18.2–32.9 | 38.8–59.1 | 28.8–39.1 | 0.046 | 0.038 |  |
| Children 3–10 years (unvaccinated) | Survey 1 (n=255) | Survey 2 (n=231) | Survey 3 (n=242) | Survey 4 (n=198) | Survey 5 (n=106) | Survey 6 (n=173) | Survey 7 (n=197) | Total (n=1402) | cPR* (95% CI) p-value | aPR* (95% CI) p-value | Relative change |
| No carriage |  |  |  |  |  |  |  |  |  |  |  |
| % (n) | 31.8 (81) | 37.7 (87) | 41.3 (100) | 62.1 (123) | 41.5 (44) | 51.5 (89) | 42.6 (84) | 43.4 (608) | - | - | - |
| 95% CI | 26.3–37.7 | 31.6–44.1 | 35.2–47.7 | 55.1–68.6 | 32.5–51.1 | 44.0–58.8 | 35.9–50.0 | 40.8–46.0 |  |  |  |
| VT |  |  |  |  |  |  |  |  |  |  |  |
| % (n) | 27.5 (70) | 21.7 (50) | 19.4 (47) | 14.1 (28) | 10.4 (11) | 11.0 (19) | 15.2 (30) | 18.2 (255) | 0.979 (0.970, 0.988) | 0.989 (0.979, 0.999) | -44.7 |
| 95% CI | 22.3–33.3 | 16.7–27.4 | 14.9–24.9 | 9.9–19.7 | 5.8–17.8 | 7.1–16.6 | 10.8–21.0 | 16.2–20.3 | <0.000 | 0.029 |  |
| NVT |  |  |  |  |  |  |  |  |  |  |  |
| % (n) | 40.8 (104) | 40.7 (94) | 39.3 (95) | 23.7 (47) | 48.1 (51) | 37.6 (65) | 42.1 (83) | 38.5 (539) | 1.000 (0.995, 1.005) | 1.005 (0.999, 1.010) | +3.2 |
| 95% CI | 34.9–46.9 | 34.5–47.2 | 33.2–45.6 | 18.3–30.2 | 38.7–57.6 | 30.6–45.0 | 35.4–49.2 | 35.9–41.0 | 0.914 | 0.101 |  |
| Adults 18-40 years (unvaccinated) | Survey 1 (n=198) | Survey 2 (n=201) | Survey 3 (n=279) | Survey 4 (n=308) | Survey 5 (n=305) | Survey 6 (n=277) | Survey 7 (n=202) | Total (n=1770) | cPR* (95% CI) p-value | aPR* (95% CI) p-value | Relative change |
| No carriage |  |  |  |  |  |  |  |  |  |  |  |
| % (n) | 60.6 (120) | 52.7 (106) | 55.6 (155) | 57.1 (176) | 61.6 (188) | 67.5 (187) | 61.4 (124) | 59.7 (1056) | - | - | - |
| 95% CI | 53.6–67.2 | 45.8–59.6 | 49.7–61.3 | 51.5–62.6 | 56.0–66.9 | 61.8–72.8 | 54.5–67.9 | 57.4–61.9 |  |  |  |
| VT |  |  |  |  |  |  |  |  |  |  |  |
| % (n) | 15.2 (30) | 14.4 (29) | 14.0 (39) | 14.3 (44) | 10.5 (32) | 9.0 (25) | 8.9 (18) | 12.3 (217) | 0.985 (0.974, 0.995) | 0.983 (0.973, 0.994) | -41.4 |
| 95% CI | 10.8–20.9 | 10.2–20.0 | 10.4–18.6 | 10.8–18.9 | 7.5–14.5 | 6.2–13.0 | 5.7–13.7 | 10.8–13.9 | 0.004 | 0.002 |  |
| NVT |  |  |  |  |  |  |  |  |  |  |  |
| % (n) | 24.2 (48) | 32.8 (66) | 30.5 (85) | 28.6 (88) | 27.9 (85) | 23.5 (65) | 29.7 (60) | 28.1 (497) | 0.997 (0.991, 1.00) | 0.997 (0.991, 1.004) | +22.7% |
| 95% CI | 18.8–30.7 | 26.7–39.6 | 25.3–36.1 | 23.8–33.9 | 23.1–33.2 | 18.8–28.8 | 23.8–36.4 | 26.0–30.2 | 0.446 | 0.392 |  |

cPR=crude prevalence ratio. aPR= adjusted prevalence ratio (adjusted for age at recruitment). CI=confidence interval. VT=vaccine serotype. NVT=non-vaccine serotype. n=total number recruited.
<sup>§</sup> There was no recruitment for these age groups during the specified surveys
<sup>¥</sup> Carriage prevalence ratios (PR) were calculated over the study duration by log-binomial regression using months (30.4 days) between study start and participant recruitment, coded as a single time variable, allowing an estimate of prevalence ratio per month.

Among HIV-infected adults on ART, aggregated (surveys 1–7) VT and NVT carriage prevalence were respectively, 12·3% (95% CI 10·8–13·9) and 28·1% (95% CI 26·0–30·2). There was a 41·4% relative reduction in VT carriage, from 15·2% (10·8–20·9) to 8·9% (5·7–13·7); aPR: 0·983 (0·973–0·994), p=0·002. There was an increase 22·7% relative increase in NVT carriage, from 24·2% (18·8–30·7) to 29·7% (23·8–36·4); aPR: 0·997 (CI 0·991–1·004), p=0·392. (Figure 2; Table 3)

Although all 13 VTs were identified in each of the three older (3 years–40 years old) study groups, with serotype 3 the predominant VT in each, serotype carriage dynamics were more heterogenous among those <3 years old. (Figure 3; Supplement 2). Serotype 1, a common cause of IPD in Africa,^38,39^ contributed 3.0% to the all-ages VT carriage prevalence.

**Figure 3.**
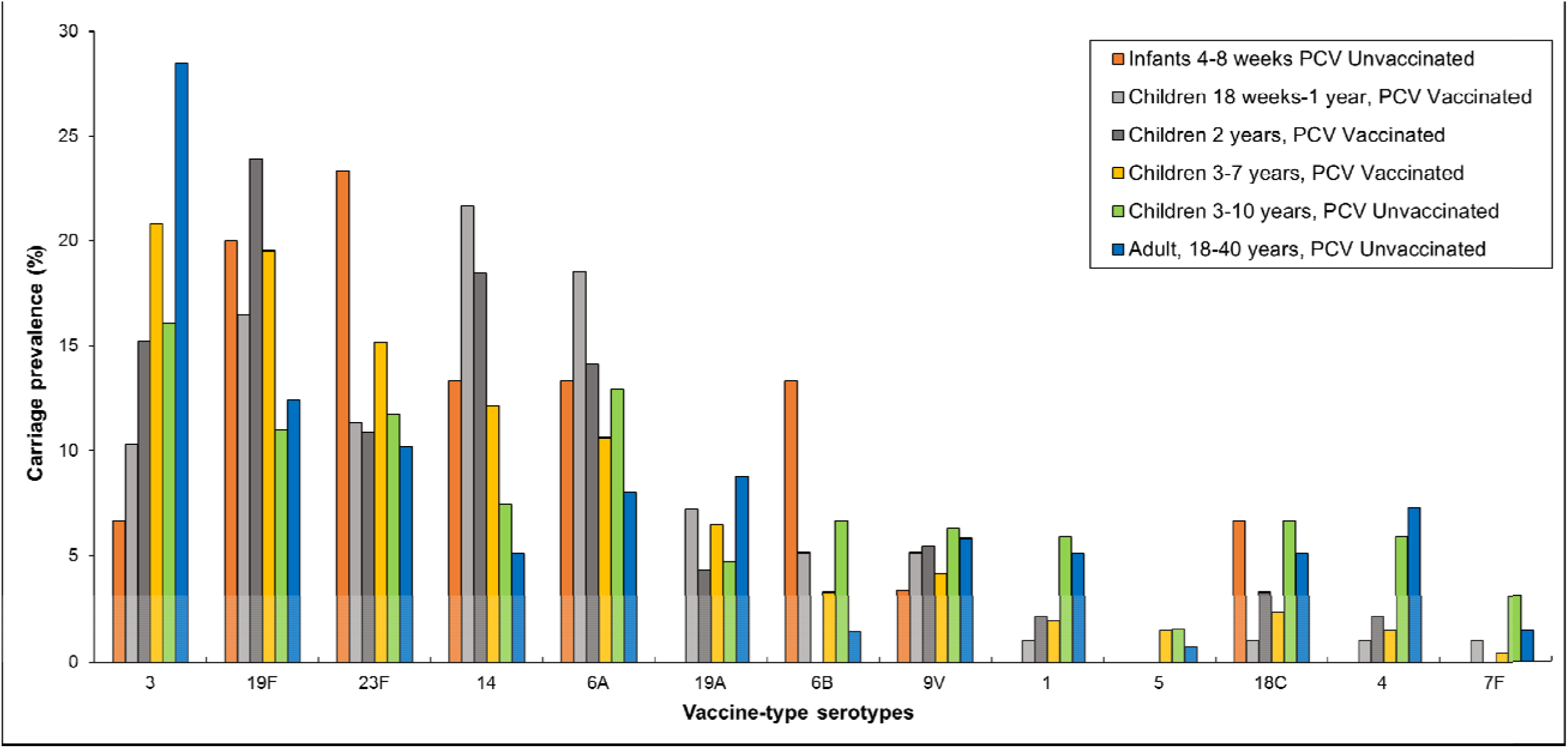
Distribution of vaccine-serotype pneumococcal carriage, aggregated across study period and stratified by study group. Proportion of vaccine-serotype (VT) carriage attributed to individual vaccine serotypes across all surveys, stratified by study group

### Probability of VT and NVT carriage with age and estimated carriage half-life

Using non-linear regression analysis to investigate the risk of VT carriage by age among children 3·6–10 years old, the probability of VT carriage was found to decline with age (Figure 4). With carriage data censored at a minimum of 3·6 years of age, the population-averaged effect of not receiving the vaccination more than doubled the probability of VT carriage, β=2·15 (95% CI 1·47–2·83) (Table 4). While the model reported different estimated probabilities of VT carriage (22% for PCV vaccinated and 47% [α*β] for PCV unvaccinated children at 3.6 years of age), the estimated half-life of VT carriage prevalence was similar (T_½_: 3·34 years [1·78–6·26] vs. T_½_: 3·26 years [2·42–4·38] respectively). Investigating probability of NVT carriage, β was 0·91 (0·73–1·09), with similar estimated probabilities of NVT carriage for vaccinated and unvaccinated children at 3.6 years of age (59% and 54% respectively [α*β]). The estimated half-life of NVT carriage prevalence was also similar among PCV-vaccinated (T_½_: 9·46 years [4·69–19·04]) and PCV-unvaccinated (T_½_: 9·83 years [5·69–16·99]) children.

**Figure 4.**
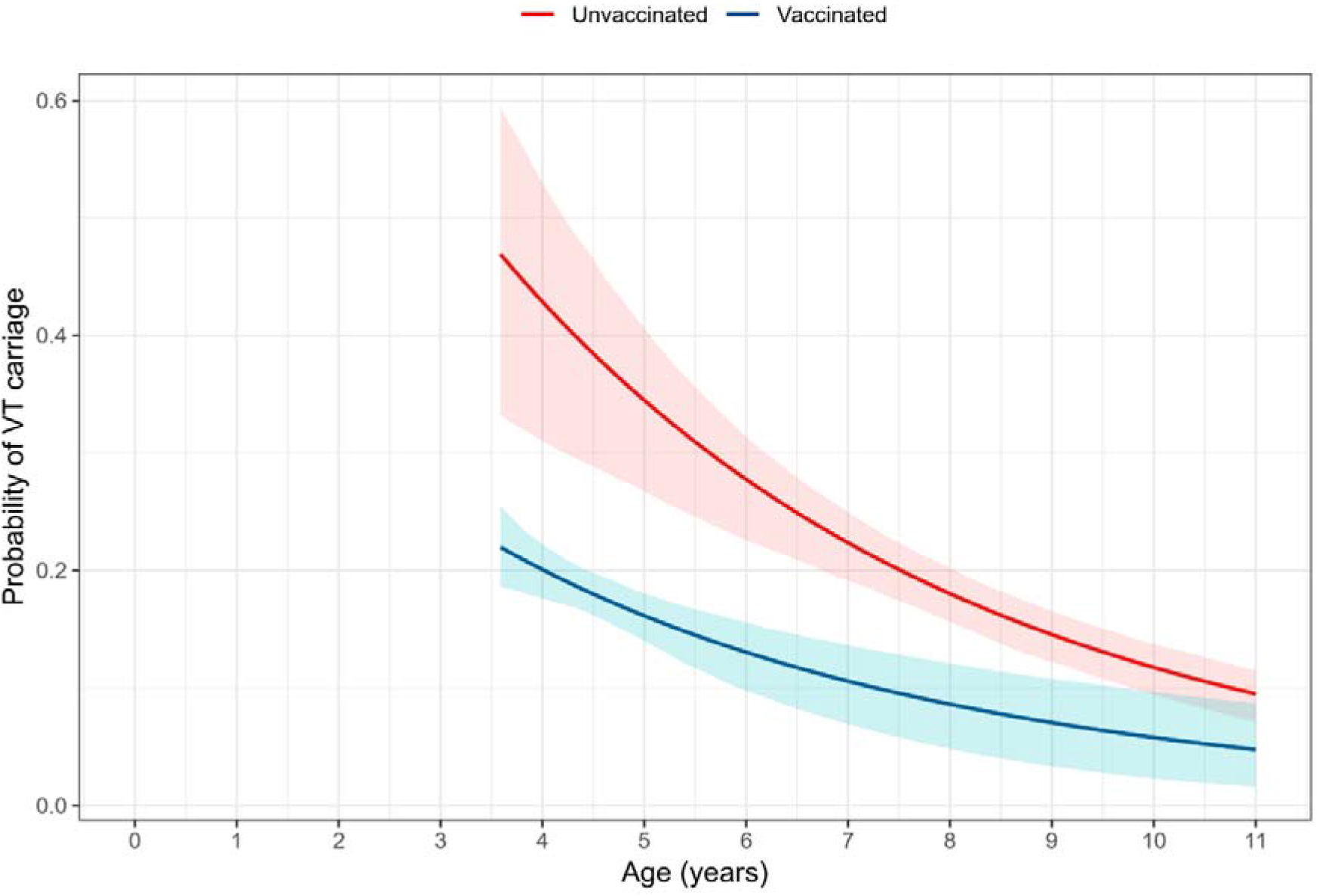
Non-linear modelling of the relationship between estimated probability of *S. pneumoniae* VT carriage and child’s age. Estimated individual probabilities and pointwise 95% confidence intervals (shaded regions) of the probability of VT carriage as a function of a child’s age (years), for an unvaccinated child (red line) and a vaccinated child (blue line). The fitted line for unvaccinated children includes the range of the empiric data. The fitted line for vaccinated children is left-censored at 3·6 years old and extrapolated beyond the oldest vaccinated child (7·9 years old). The model shows significantly different estimated probabilities of VT carriage, while the half-life of VT carriage translates to very similar estimates among PCV-vaccinated (3·34 years) and PCV-unvaccinated (3·26 years) children.

**Table 4:**
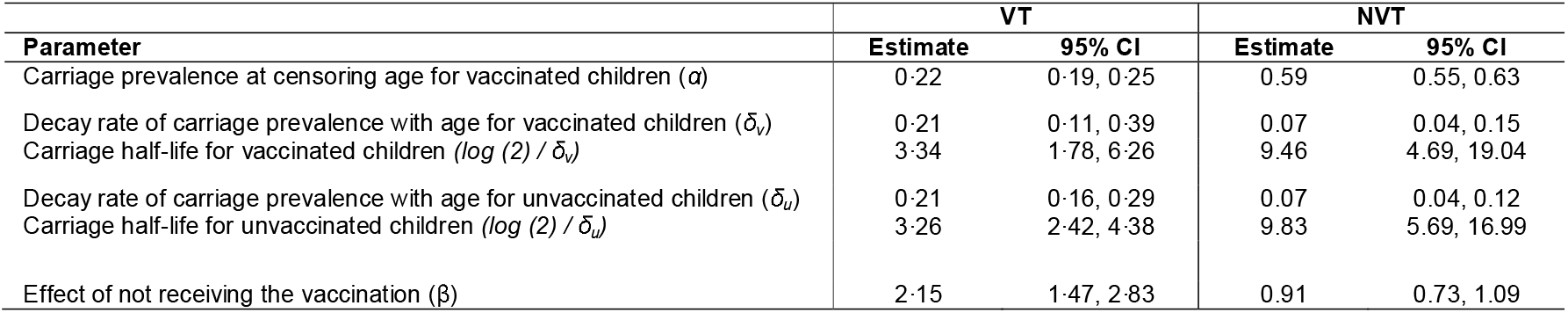
Maximum likelihood estimates for the fitted model for probability of carriage with age and estimated carriage half-life, censored at 3.6 years of age

| Parameter | VT |  | NVT |  |
| --- | --- | --- | --- | --- |
|  | Estimate | 95% CI | Estimate | 95% CI |
| Carriage prevalence at censoring age for vaccinated children ($\alpha$) | 0.22 | 0.19, 0.25 | 0.59 | 0.55, 0.63 |
| Decay rate of carriage prevalence with age for vaccinated children ($\delta_v$) | 0.21 | 0.11, 0.39 | 0.07 | 0.04, 0.15 |
| Carriage half-life for vaccinated children ($\log(2) / \delta_v$) | 3.34 | 1.78, 6.26 | 9.46 | 4.69, 19.04 |
| Decay rate of carriage prevalence with age for unvaccinated children ($\delta_u$) | 0.21 | 0.16, 0.29 | 0.07 | 0.04, 0.12 |
| Carriage half-life for unvaccinated children ($\log(2) / \delta_u$) | 3.26 | 2.42, 4.38 | 9.83 | 5.69, 16.99 |
| Effect of not receiving the vaccination ($\beta$) | 2.15 | 1.47, 2.83 | 0.91 | 0.73, 1.09 |

Assessment of the goodness-of-fit indicates a good fit, with no discernible relationship between the residual and the predicted values and the range of residuals compatible with the theoretical mean and the standard deviations of 0 and 1, respectively (Supplement 1, Figure 3).

## DISCUSSION

In this community-based assessment of pneumococcal carriage we surveyed potential reservoir populations after the introduction of routine PCV13 in Malawi. Over the 3.5-year study period, we found only a modest relative decline in VT carriage among recently vaccinated children 18 weeks to 1 year old (21%), vaccinated children 2 years old (31%) and vaccinated children 3–7 years old (16%). All 13 VTs were isolated, despite high vaccine uptake and good schedule adherence. There was only a modest increase in NVT carriage. The 18% residual aggregated VT carriage prevalence among PCV-vaccinated children was consistent with the 16.5% previously reported among PCV-vaccinated children 1-4 years old in northern Malawi, 3.5 years after PCV13 introduction.^11^ Though these reported residual VT carriage prevalences were lower than that observed in northern Malawi before vaccine introduction (28%),^11^ they did not reach the substantially lower levels rapidly achieved in high-income low-carriage prevalence settings (<5%) associated with control of carriage and transmission. ^22–24^ VT carriage dynamics did not conform to a simple exponential distribution, with slightly higher VT carriage prevalence among PCV-vaccinated children under 4 years of age than older vaccinated children. The trend of reduced VT carriage was also evident in older unvaccinated children, suggesting that the effect is age-driven and not simply vaccine dependent (Supplement 1). This underlines the complex relationship in the first few years of life between VT carriage and the impact of waning vaccine-induced mucosal immunity and acquisition of natural immunity. We found a more marked decline in VT carriage among unvaccinated (age-ineligible) children 3–10 years old (45%), and HIV-infected adults on ART (41%). In the light of the recent WHO Technical Expert Consultation Report on Optimization of PCV Impact,^40^ these data start to address the paucity of information on the long-term impact of the widely implemented 3+0 vaccine schedules on serotype-specific disease and carriage in this region.

The lower probability of carriage among vaccinated children suggests vaccination does reduce the probability of VT carriage, providing a lower VT carriage setpoint. However, much of that direct vaccine-induced protection occurs quite early, perhaps within the first 6 months of life, and we suggest that thereafter the decline in carriage probability is determined by natural immunity. Indeed, our non-linear statistical analysis shows comparable half-life of VT carriage between PCV-vaccinated and PCV-unvaccinated children beyond the age of 3.6 years. The mechanism underlying the vaccine effect could be prevention of carriage (reduced incidence) or shortening of carriage duration (reduced point prevalence). We further postulate that in older vaccinated and unvaccinated children, the reductions in carriage prevalence are due to the indirect benefits of vaccination augmented by naturally acquired immunity to subcapsular protein antigens.^41,42^

To achieve herd protection in settings with high carriage prevalence, such as Malawi, we need to effectively interrupt person-to-person transmission. In Finland, a microsimulation model suggested a moderate transmission potential of pneumococcal carriage, predicting the elimination of VT carriage among those vaccinated within 5–10 years of PCV introduction, assuming high (90%) vaccine coverage and moderate (50%) vaccine efficacy against acquisition.^43^ Thus, vaccine impact predicted by transmission models from low carriage prevalence settings probably does not translate to high carriage prevalence settings. Although it has previously been assumed that PCVs would eliminate VT carriage in mature PCV programmes,^44^ our data bring into question the potential for either a sustained direct or indirect effect on carriage using a 3+0 strategy.

In Malawi, the vaccine impact on carriage prevalence has been less than that observed in Kenya, The Gambia and South Africa which have used different vaccination strategies. Kenya reported a reduction from 34% to 9% VT carriage among PCV-vaccinated children under 5 years of age, 6 years after introduction of 10-valent PCV.^19^ The Gambia reported a reduction from 50% to 13% VT carriage among children 2–5 years old, 20 months after introducing the 7-valent PCV.^45^ Likewise, a study from South Africa showed reduced PCV13-serotype colonisation from 37% to 13% within 1 year of transitioning from PCV7 to PCV13.^46^ However, these countries have also not achieved the low carriage prevalences seen in Europe and North America 2 to 3 years post introduction.^3,47^ We propose that a high force of infection (FOI) in settings such as Malawi limits a 3+0 schedule to achieving only a short duration of VT carriage control in infants. While a 2+1 schedule, as deployed in South Africa, may improve colonisation control, this remains unproven in other African settings. Given the likely importance of an early reduction in transmission intensity to maintain a reduced carriage prevalence, a catch-up-campaign with booster doses over a broader age range (i.e. <5 years of age) may also be required. Although the Global Alliance for Vaccines and Immunization (GAVI) has considerably reduced PCV costs for low-income countries,^48,49^ vaccine impact must be optimised (particularly indirect effects) to achieve financial sustainability. The FOI and determinants of transmission between and within age groups need to be considered, as new approaches to improving vaccine-induced carriage reduction are proposed and tested.

Unlike low-transmission settings,^50^ as well as The Gambia^25^ and South Africa,^46^ we observed a modest increase in NVT carriage among children in Malawi. Given evidence elsewhere of rapid serotype replacement after PCV introduction, it is possible that serotype replacement and redistribution had already occurred before the start of this study, and that as part of a stochastic secular trend, we are now observing an overall decrease in pneumococcal carriage prevalence. There may have also been individual NVT that increased, while other NVT decreased, in prevalence. Though distribution of individual NVT serotypes warrant further analysis, our latex serotyping methods did not allow for identifying individual NVT serotypes. It is also plausible that overall improvement in living conditions (improved nutrition, sanitation and disease control) and health care (antiretroviral roll-out and rotavirus vaccination) have allowed a sustained drop in pneumococcal carriage as a result of improved health, evidenced by falling under 5 mortality in recent years.^51^ Either way, the importance of these trends in NVT carriage will become clearer as the trends in NVT invasive disease become available from these different settings.

We have previously shown incomplete pneumococcal protein antigen-specific reconstitution of natural immunity and high levels of pneumococcal colonisation in HIV-infected Malawian adults on ART.^33^ We now show that the adult population has not greatly benefitted from indirect protection against carriage following routine infant PVC13 introduction and indeed may represent a reservoir of VT carriage and transmission. Previous studies in Malawi and South Africa have suggested that despite a higher risk of VT pneumococcal colonisation among HIV-infected women, they are still unlikely to be a significant source of transmission to their children.^14,52^ However, in the context of routine infant PCV13 and rapid waning of vaccine-induced immunity, the balance of transmission may now be different. Given the higher risk of IPD, ongoing burden of pneumococcal pneumonia,^53,54^ and the evidence that PCV protects HIV-infected adults from recurrent VT pneumococcal infections,^55^ targeted vaccination benefitting this at-risk population may help reduce overall carriage and disease prevalence.

### Limitations

This work provides a robust community-based estimate of VT and NVT pneumococcal carriage in Blantyre. The study was conducted over a relatively short timeframe for understanding long-term temporal trends. For this reason, the statistical analysis is limited in its ability to disentangle the effects of calendar time and age-since-vaccination, given the small overlap in ages of vaccinated and unvaccinated children in our data. Although there are pre-vaccine-introduction data from elsewhere in Malawi, there are no equivalent historical carriage data for urban settings in Malawi using the same sampling frame. However, this does not detract from the finding of high levels of residual VT carriage in these reservoir populations. Finally, given evidence that more sensitive serotyping methods that detect multiple serotype carriage (e.g. DNA microarray) will increase VT carriage estimates, our carriage prevalence data likely underestimate the true residual VT prevalence levels.^56–58^

## CONCLUSION

Despite success in achieving direct protection of infants against disease, a 3+0 PCV13 schedule in Malawi has not achieved the low universal VT carriage prevalence reported in high-income settings and that is required to control carriage and transmission. We propose that although vaccine-induced immunity reduces the risk of VT carriage in children up to approximately 6 months of age, in the context of a high residual FOI, this impact is limited by rapid waning of vaccine-induced mucosal immunity and pneumococcal recolonisation (Figure 5). Therefore, alternative schedules and vaccine introduction approaches in high pneumococcal carriage, high-disease-burden countries should be revisited through robust evaluation rather than through programmatic change without supporting evidence. Furthermore, we need to better understand the relative impact of waning vaccine-induced immunity, indirect vaccine protection and naturally-acquired immunity on VT carriage in the two to three years after vaccination.

**Figure 5.**
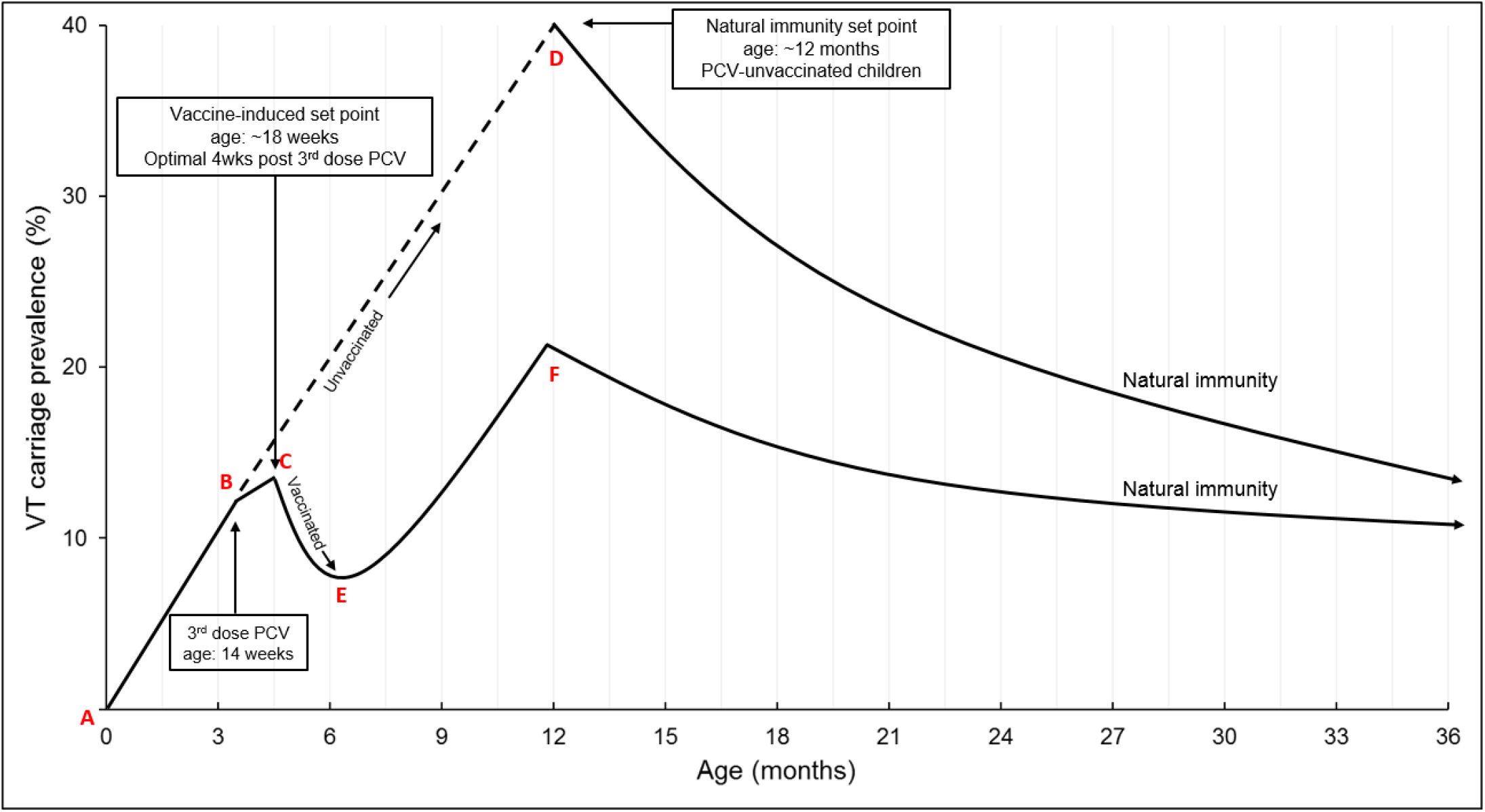
A hypothesis – the role of pneumococcal conjugate vaccine-induced and natural anti-pneumococcal immunity in determining the prevalence of colonisation by *S. pneumoniae* among children in a setting with high carriage prevalence. **A:** Children born uncolonised with *S. pneumoniae*. **A–B:** Soon after birth, children are colonised with VT and NVT pneumococcus via contact with family and community members. **B:** At 14 weeks of age, vaccine-eligible children have received 3 doses PCV13, with an optimal immunogenic response for vaccine-induced mucosal immunity 4 weeks later (**C**). Among PCV-vaccinated children, 18-weeks is the approximate vaccine-induced set point, with a rapid decrease in VT prevalence (**C–E**) until 6 months of age (**E**). At 6 months of age, there is an increase in risk of VT carriage (**E–F**), driven by increased force of infection in the context of waning vaccine-induced immunity, the former due partly to increased contact with other young children in the household and community. VT carriage prevalence increases until naturally acquired immunity starts to impact on colonisation (**F**), reducing pneumococcal carriage prevalence. Among PCV-unvaccinated children, risk of VT carriage continues to increase largely unchecked (**B–D**) until naturally acquired immunity starts to impact on colonisation (**D**), reducing pneumococcal carriage prevalence. Among these unvaccinated children, 12 months is the approximate set-point induced by naturally-acquired immunity. Indirect vaccine effects will impact on the height of C–E–F and B–D, as well as the rate of decline in VT carriage prevalence.

## Acknowledgements

We thank the individuals who participated in this study and the local schools and authorities for their support. We are grateful to the study field teams (supported by Farouck Bonomali and Roseline Nyirenda) and the study laboratory team. We are grateful to the hospitality of the QECH ART Clinic, led by Ken Malisita. Our thanks also extend to the MLW laboratory management team (led by Brigitte Denis) and the MLW data management team (led by Clemens Masesa). RSH, NF and TS are supported by the National Institute for Health Research (NIHR) Global Health Research Unit on Mucosal Pathogens using UK aid from the UK Government. The views expressed in this publication are those of the author(s) and not necessarily those of the NIHR or the Department of Health and Social Care”.

## Author contributions

T.D.S., N.B.Z., D.E., N.F., and R.S.H. designed the study. All contributed to the development or design of methodology. T.D.S., N.F. and R.H.S. oversaw the study, data collection, and data management. C.F. and P.D. developed the statistical regression analysis. T.D.S., N.F., and R.S.H. conducted the statistical analysis. T.D.S., N.F., and R.S.H. wrote the first draft of the paper, and all authors contributed to subsequent drafts. All read and approved the final version of the report.

## Competing interests

Dr. Bar-Zeev reports investigator-initiated research grants from GlaxoSmithKline Biologicals and from Takeda Pharmaceuticals outside the submitted work. No other competing interests were reported by authors.

## Materials & Correspondence

Correspondence and material requests should be addressed to: Todd Swarthout, Malawi-Liverpool-Wellcome Trust Clinical Research Programme, P.O. Box 30096, Chichiri Blantyre 3, Malawi;

## Data availability

The data that support the findings of this study are available from the corresponding author upon reasonable request.

## Funding

Bill & Melinda Gates Foundation, Wellcome Trust UK, Medical Research Council, NIHR. The funders had no role in study design, collection, analysis, data interpretation, writing of the report or in the decision to submit the paper for publication. The corresponding author had full access to the study data and, together with the senior authors, had final responsibility for the decision to submit for publication.

## Ethics approval and consent to participate

The study protocol was approved by the College of Medicine Research and Ethics Committee, University of Malawi (P.02/15/1677) and the Liverpool School of Tropical Medicine Research Ethics Committee (14.056). Adult participants and parents/guardians of child participants provided written informed consent, children 8-10 years old provided informed assent. This included consent for publication.

## Notes

#### Summary of Updates

The previous version included analysis from approximately 2 years of pneumococcal carriage surveillance from children 3-10 years of age and HIV-infected adults on ART. This revised version has an expanded analysis that includes data from a total of approximately 3.5 years of pneumococcal carriage surveillance. This includes pneumococcal carriage data from data included in our first version, as well as children 4-8 weeks of age (prior to first dose PCV) add PCV-vaccinated children 18 weeks to 2 years of age. These younger age groups were recruited starting approximately 1.5 years after study start. The non-linear model analysis, used to assess risk of VT carriage by age, was expanded to included risk of NVT carriage by age.

