## Supplement 1 Figure 1; Supplement 1 Figure 2; Supplement 1 Figure 3; Supplement 2a, Table 1; Supplement 2b, Table 1; Supplement 3, Table 1 for "High residual prevalence of vaccine-serotype *Streptococcus pneumoniae* carriage after introduction of a pneumococcal conjugate vaccine in Malawi: a prospective serial cross-sectional study"

### Supplement 1: Non-linear regression framework, goodness-of-fit, limitations and statistical details

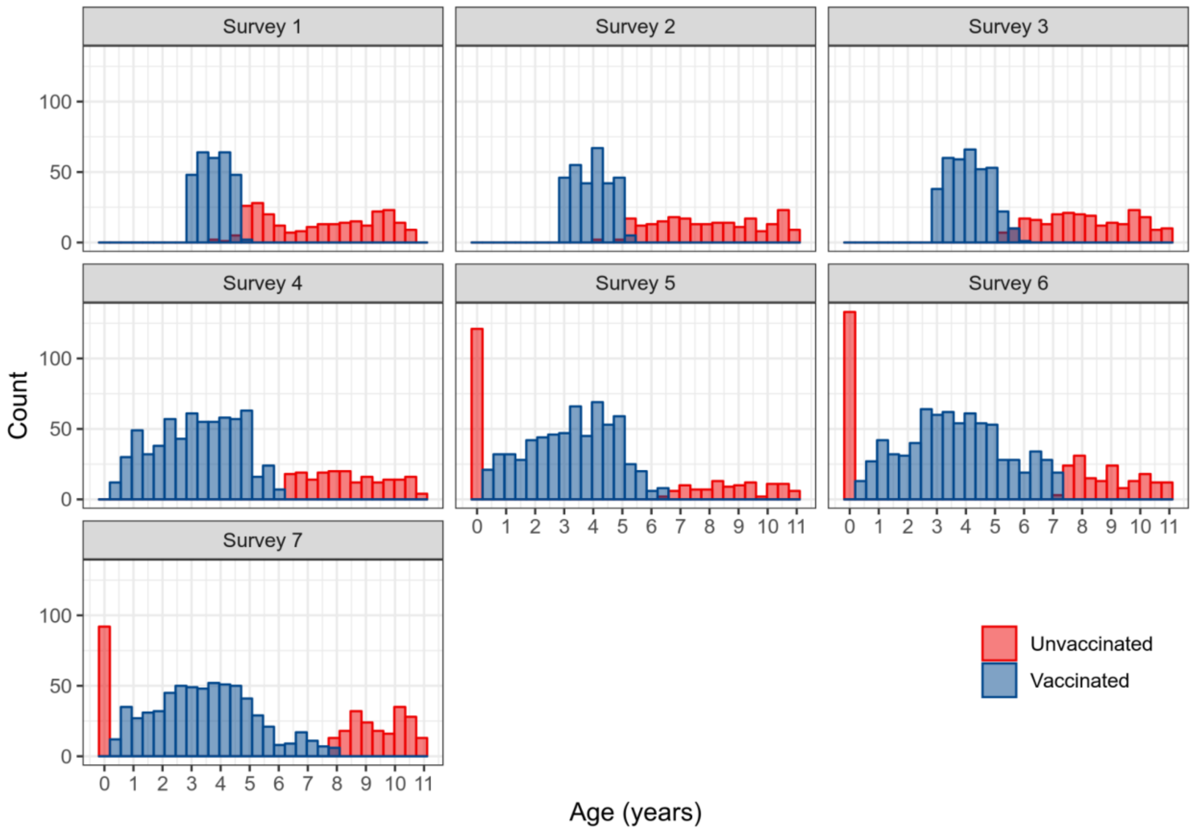


**Figure 1. Age distribution of vaccinated and unvaccinated children stratified by survey.**

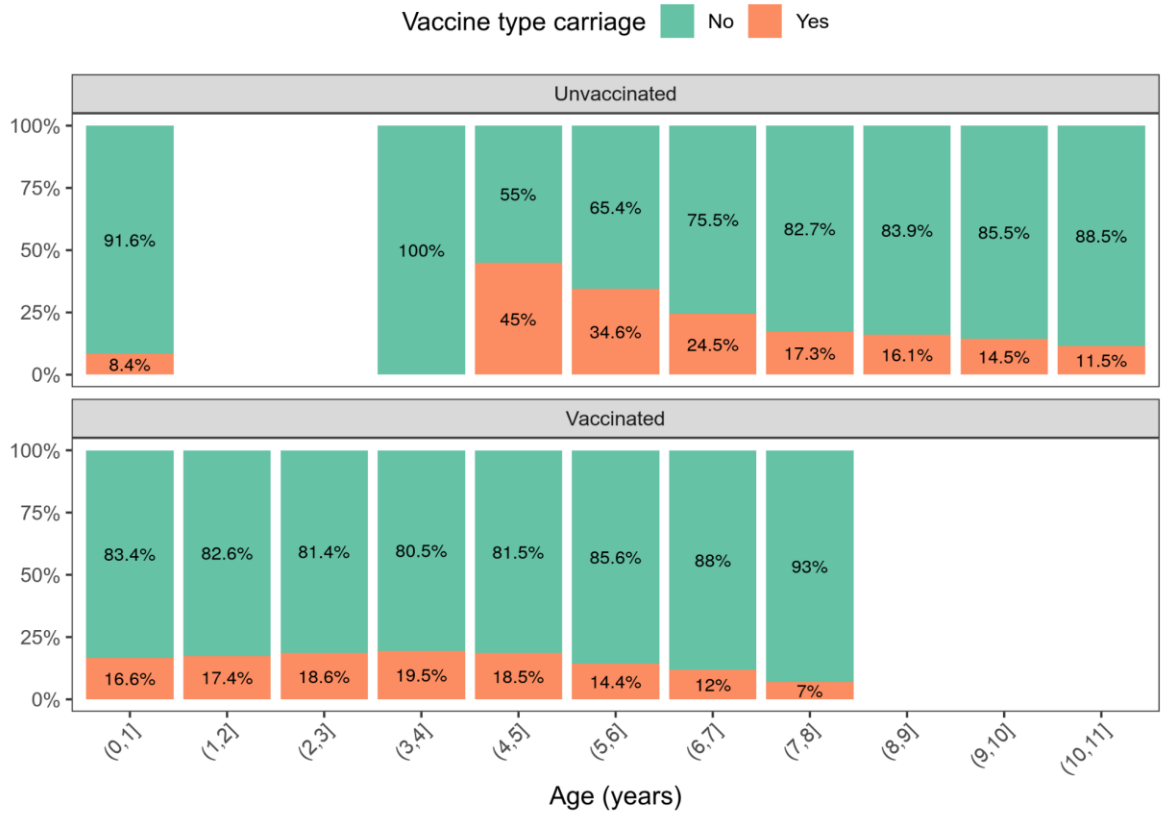

**Figure 2. Proportion of vaccine type carriage by age group in vaccinated and unvaccinated children.**

### Model specification, VT

In this cross-sectional study, the response for each child is a single binary variable, $Y_i = 1/0$ representing presence/absence of VT carriage at the time of measurement, with respective probabilities $p_i$ and $1 - p_i$. The modelled probability, $p_i$, of VT carriage for child $i$ is $\alpha\beta\exp\{-\delta_u(age_i - t_c)\}$ if the child is unvaccinated and $\alpha\exp\{-\delta_v(age_i - t_c)\}$ if vaccinated, where $age_i$ is the age in years of child $i$ at time of measurement, $t_c$ is the time at which we censor the data (3.6 years), $\beta$ is the effect of not receiving the vaccination, $\delta_u$ and $\delta_v$ are the rates of decay of VT carriage prevalence with age for unvaccinated and vaccinated children respectively and $\alpha$ is the prevalence for unvaccinated children at time $t_c$.

### Goodness-of-fit and limitations

The conventional way to assess goodness-of-fit of a regression model is to plot standardised residuals against predicted (fitted) values; in a well-fitting model, the plot should show no discernible structure. However, individual residuals from a model with binary response are unreliable. Instead, we use grouped residuals as follows. Within each of the unvaccinated and vaccinated sets of children, order the estimated probabilities $p_i$ from smallest to largest. Choose a group size, $k$. The first grouped predicted value is the sum $s_1 = p_1 + \cdots + p_k$, and the first standardised grouped residual is $r_1 = \{(Y_1 + \cdots + Y_k) - s_1\}/\sqrt{(v_1)}$ where $Y_i = 0/1$ denotes absence/presence of VT carriage in the ith child and $v_1 = p_1(1 - p_1) + \cdots + p_k(1 - p_k)$. The second grouped predicted value and standardised residual are calculated in the same way using the next $k$ ordered $p_i$, and so on.

Figure 3 shows the resulting plot, using $k = 125$ and $k = 93$ for unvaccinated and vaccinated children, respectively, to give 15 groups in each of the two sets. The plot indicates a good fit in that (a) there is no discernible relationship between the residual and predicted values, and (b) the range of the residuals, from approximately -2 to +2, is compatible with their theoretical mean and standard deviation of 0 and 1, respectively, if the model is correct.

The major limitation of the analysis reported here is the inability of our model to extrapolate before the censoring time $t_c$ with a reasonable amount of uncertainty. This is due to the small overlap in the age ranges of vaccinated and unvaccinated children and to the fact that the VT carriage dynamic for vaccinated children is too complex to capture, with the available data, in the early years of life.

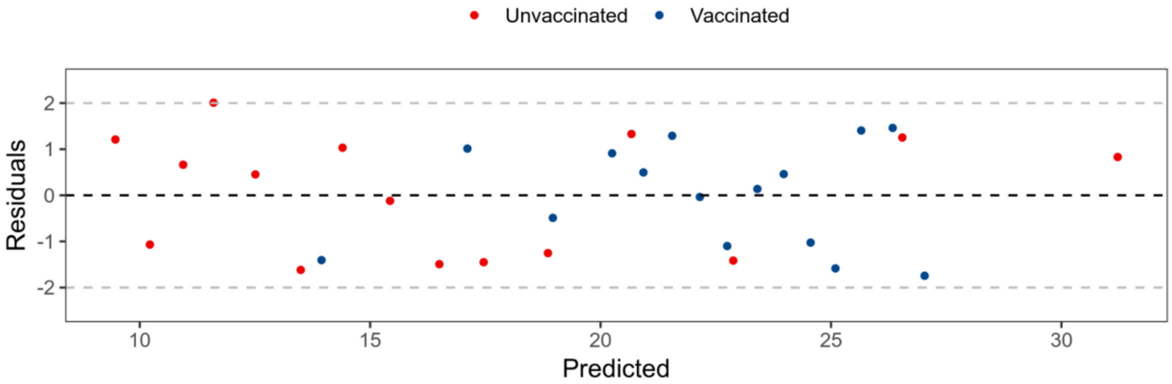


**Figure 3. Plot of standardised grouped residuals against predicted values; see text for detailed explanation.**

**Supplement 2a, Table 1. Proportion and frequency of VT carriage attributed to each VT, stratified by study group & survey, PCV-vaccinated children**

| VT serotype | Survey-1 % (n) | Survey-2 % (n) | Survey-3 % (n) | Survey-4 % (n) | Survey-5 % (n) | Survey-6 % (n) | Survey-7 % (n) | Total % (n) |
|---|---|---|---|---|---|---|---|---|
| **Children 18 weeks to 1 year *** | | | | | | | | |
| 1 | - | - | - | 0 | 0 | (4.4) 1 | 0 | 1.0 (1) |
| 3 | - | - | - | 6.9 (2) | 7.7 (2) | 8.7 (2) | 21.0 (4) | 10.3 (10) |
| 4 | - | - | - | 0 | 0 | 0 | 5.3 (1) | 1.0 (1) |
| 5 | - | - | - | 0 | 0 | 0 | 0 | 0 |
| 6A | - | - | - | 20.7 (6) | 15.4 (4) | 21.7 (5) | 15.8 (3) | 18.6 (18) |
| 6B | - | - | - | 13.8 (4) | 3.9 (1) | 0 | 0 | 5.2 (5) |
| 7F | - | - | - | 0 | 3.9 (1) | 0 | 0 | 1.0 (1) |
| 9V | - | - | - | 3.5 (1) | 7.7 (2) | 4.4 (1) | 5.3 (`) | 5.2 (5) |
| 14 | - | - | - | 13.8 (4) | 15.4 (4) | 34.8 (8) | 26.3 (5) | 21.7 (21) |
| 18C | - | - | - | 0 | 0 | 0 | 5.3 (1) | 1.0 (1) |
| 19A | - | - | - | 13.8 (4) | 3.9 (1) | 8.7 (2) | 0 | 7,2 (7) |
| 19F | - | - | - | 13.8 (4) | 23.1 (6) | 17.4 (4) | 10.5 (2) | 16.5 (16) |
| 23F | - | - | - | 13.8 (4) | 19.2 (5) | 0 | 10.5 (2) | 11.3 (11) |
| Total | - | - | - | 29 | 26 | 23 | 19 | 97 |
| **Children 2 years (PCV vaccinated) *** | | | | | | | | |
| 1 | - | - | - | 0 | (2) | 0 | 0 | 2.2 (2) |
| 3 | - | - | - | 18.5 (5) | 14.3 (3) | 16.0 (4) | 10.5 (2) | 15.2 (14) |
| 4 | - | - | - | 0 | 0 | 0 | 10.5 (2) | 2.2 (2) |
| 5 | - | - | - | () | () | () | () | () |
| 6A | - | - | - | 22.2 (6) | 9.5 (2) | 8.0 (2) | 15.8 (3) | 14.1 (13) |
| 6B | - | - | - | () | () | () | () | () |
| 7F | - | - | - | () | () | () | () | () |
| 9V | - | - | - | 7.4 (2) | 9.5 (2) | 0 | 5.3 (1) | 5.4 (5) |
| 14 | - | - | - | 11.1 (3) | 14.3 (3) | 28.0 (7) | 21.1 (4) | 18.5 (17) |
| 18C | - | - | - | 3.7 (1) | 4.8 (1) | 4.0 (1) | 0 | 3.3 (3) |
| 19A | - | - | - | 7.4 (2) | 0 | 4.0 (1) | 5.3 (1) | 4.4 (4) |
| 19F | - | - | - | 25.9 (7) | 14.3 (3) | 28.0 (7) | 26.3 (5) | 23.9 (22) |
| 23F | - | - | - | 3.7 (1) | 23.8 (5) | 12.0 (3) | 5.3 (1) | 10.9 (10) |
| Total | | | | (100) 27 | (100) 21 | (100) 25 | (100) 19 | (100) 92 |
| **Children 3-7 years (PCV vaccinated)** | | | | | | | | |
| 1 | 5.3 (3) | 1.6 (1) | 1.3 (1) | 0 | 4.1 (3) | 1.6 (1) | 0 | 2.0 (9) |
| 3 | 17.5 (10) | 22.6 (14) | 25.3 (19) | 17.9 (12) | 24.3 (18) | 11.1 (7) | 25.4 (16) | 20.8 (96) |
| 4 | 0 | 3.2 (2) | 1.3 (1) | 0 | 0 | 6.4 (4) | (0) | 1.5 (7) |
| 5 | 3.5 (2) | 4.8 (3) | 0 | 1.5 (1) | 0 | 1.6 (1) | 0 | 1.5 (7) |
| 6A | 14.0 (8) | 6.5 (4) | 12.0 (9) | 9.0 (6) | 9.5 (7) | 17.5 (11) | 6.4 (4) | 10.6 (49) |
| 6B | 1.8 (1) | 4.8 (3) | 6.7 (5) | 1.5 (1) | 2.7 (2) | 1.6 (1) | 3.2 (2) | 3.3 (15) |
| 7F | 0 | 0 | 0 | 1.5 (1) | 0 | 1.6 (1) | 0 | 0.4 (2) |
| 9V | 3.5 (2) | 6.5 (4) | 1.3 (1) | 3.0 (2) | 4.1 (3) | 4.8 (3) | 6.4 (4) | 4.1 (19) |
| 14 | 8.8 (5) | 11.3 (7) | 6.7 (5) | 14.9 (10) | 10.8 (8) | 14.3 (9) | 19.1 (12) | 12.2 (56) |
| 18C | 1.8 (1) | 0 | 4.0 (3) | 3.0 (2) | 1.4 (1) | 3.2 (2) | 3.2 (2) | 2.4 (11) |
| 19A | 7.0 (4) | 12.9 (8) | 9.3 (7) | 6.0 (4) | 6.8 (5) | 1.6 (1) | 1.6 (1) | 6.5 (30) |
| 19F | 24.6 (14) | 8.1 (5) | 21.3 (16) | 28.4 (19) | 13.5 (10) | 17.5 (11) | 23.8 (15) | 19.5 (90) |
| 23F | 12.3 (7) | 17.7 (11) | 10.7 (8) | 13.4 (9) | 23.0 (17) | 17.5 (11) | 11.1 (7) | 15.2 (70) |
| Total | 57 | 62 | 75 | 67 | 74 | 63 | 63 | 461 |

*Survey 1-3: Data not collected

**Supplement 2b, Table 1. Proportion and frequency of VT carriage attributed to each VT, stratified by study group & survey, PCV-unvaccinated children**

| VT serotype | Survey-1 % (n) | Survey-2 % (n) | Survey-3 % (n) | Survey-4 % (n) | Survey-5 % (n) | Survey-6 % (n) | Survey-7 % (n) | Total % (n) |
|---|---|---|---|---|---|---|---|---|
| **Children 4-8 weeks *** | | | | | | | | |
| 1 | - | - | - | - | 0 | 0 | 0 | 0 |
| 3 | - | - | - | - | 7.7 (1) | 0 | 12.5 (1) | 6.7 (2) |
| 4 | - | - | - | - | 0 | 0 | 0 | 0 |
| 5 | - | - | - | - | 0 | 0 | 0 | 0 |
| 6A | - | - | - | - | 15.4 (2) | 25.0 (2) | 0 | 4 (13.3) |
| 6B | - | - | - | - | 7.7 (1) | 0 | 37.5 (3) | 4 (13.3) |
| 7F | - | - | - | - | 0 | 0 | 0 | 0 |
| 9V | - | - | - | - | 1 (7.7) | 0 | 0 | 1 (3.3) |
| 14 | - | - | - | - | 7.7 (1) | 25.0 (2) | 12.5 (1) | 13.3 (4) |
| 18C | - | - | - | - | 7.7 (1) | 12.5 (1) | 0 | 6.7 (2) |
| 19A | - | - | - | - | 0 | 0 | 0 | 0 |
| 19F | - | - | - | - | 15.4 (2) | 25.0 (2) | 25.0 (2) | 20.0 (6) |
| 23F | - | - | - | - | 30.8 (4) | 12.5 (1) | 12.5 (1) | 23.3 (6) |
| **Total** | - | - | - | - | 13 | 8 | 8 | 29 |
| **Children 3-10 years** | | | | | | | | |
| 1 | 14.3 (10) | 6.0 (3) | 2.1 (1) | 0 | 0 | 5.3 (1) | 0 | 5.6 (15) |
| 3 | 12.9 (9) | 20.0 (10) | 23.4 (11) | 14.3 (4) | 0 | 26.3 (5) | 6.7 (2) | 16.1 (41) |
| 4 | 5.7 (4) | 2.0 (1) | 10.6 (5) | 7.1 (2) | 0 | 15.8 (3) | 0 | 5.9 (15) |
| 5 | 1.4 (1) | 0 | 0 | 0 | 0 | 10.5 (2) | 3.3 (1) | 1.6 (4) |
| 6A | 10,0 (7) | 20.0 (10) | 6.4 (3) | 21.4 (6) | 18.2 (2) | 5.3 (1) | 13.3 (4) | 12.9 (33) |
| 6B | 2.9 (2) | 10.0 (5) | 8.5 (4) | 7.1 (2) | 9.1 (1) | 0 | 10.0 (3) | 6.7 (17) |
| 7F | 5.7 (4) | 2.0 (1) | 2.1 (1) | 0 | 9.1 (1) | 5.3 (1) | 0 | 6.7 (8) |
| 9V | 8.6 (6) | 6.0 (3) | 4.3 (2) | 7.1 (2) | 9.1 (1) | 0 | 6.7 (2) | 6.3 (16) |
| 14 | 0 | 8.0 (4) | 17.0 (8) | 7.1 (2) | 9.1 (1) | 5.3 (1) | 10.0 (3) | 7.5 (19) |
| 18C | 7.1 (5) | 4.0 (2) | 8.5 (4) | 0 | 27.3 (3) | 5.3 (1) | 6.7 (2) | 6.7 (17) |
| 19A | 4.3 (3) | 4.0 (2) | 4.3 (2) | 7.1 (2) | 0 | 0 | 10.0 (3) | 4.7 (12) |
| 19F | 12.9 (9) | 10.0 (5) | 8.5 (4) | 10.7 (3) | 0 | 10.5 (2) | 16.7 (5) | 11.0 (28) |
| 23F | 14.3 (10) | 8.0 (4) | 4.3 (2) | 17.9 (5) | 18.2 (2) | 10.5 (2) | 16.7 (5) | 11.8 (30) |
| Total | 100 (70) | 100 (50) | 100 (47) | 100 (28) | 100 (11) | 100 (19) | 100 (30) | 100 (255) |
| **Adults 18-40 years, HIV-infected on ART** | | | | | | | | |
| 1 | 3.3 (1) | 13.8 (4) | 7.7 (3) | 0 | 0 | 0 | 0 | 3.7 (8) |
| 3 | 23.3 (7) | 34.5 (10) | 28.2 (11) | 27.3 (12) | 37.5 (12) | 36.0 (9) | 55.6 (10) | 32.7 (71) |
| 4 | 13.3 (4) | 6.9 (2) | 7.7 (3) | 2.3 (1) | 9.4 (3) | 8.0 (2) | 0 | 6.9 (15) |
| 5 | 0 | 0 | 2.6 (1) | 0 | 0 | 0 | 0 | 0.5 (1) |
| 6A | 13.3 (4) | 3.5 (1) | 10.3 (4) | 4.6 (2) | 6.3 (2) | 4.0 (1) | 5.6 (1) | 6.9 (15) |
| 6B | 3.3 (1) | 3.5 (1) | 0 | 0 | 3.1 (1) | 4.0 (1) | 0 | 1.8 (4) |
| 7F | 0 | 0 | 0 | 4.6 (2) | 0 | 0 | 0 | 0.9 (2) |
| 9V | 3.3 (1) | 3.5 (1) | 5.1 (2) | 9.1 (4) | 3.1 (1) | 4.0 (1) | 0 | 4.6 (10) |
| 14 | 0 | 0 | 7.7 (3) | 9.1 (4) | 3.1 (1) | 8.0 (2) | 0 | 4.6 (10) |
| 18C | 6.7 (2) | 3.5 (1) | 0 | 9.1 (4) | 0 | 8.0 (2) | 16.7 (3) | 5.5 (12) |
| 19A | 10.0 (3) | 6.9 (2) | 7.7 (3) | 9.1 (4) | 9.4 (3) | 12.0 (3) | 5.6 (1) | 8.8 (19) |
| 19F | 20.0 (6) | 17.2 (5) | 5.1 (2) | 13.6 (6) | 15.6 (5) | 8.0 (2) | 16.7 (3) | 13.4 (29) |
| 23F | 3.3 (1) | 6.9 (2) | 18.0 (7) | 11.4 (5) | 12.5 (4) | 8.0 (2) | 0 | 9.7 (21) |
| Total | 100 (30) | 100 (29) | 100 (39) | 100 (44) | 100 (32) | 100 (25) | 100 (18) | 100 (217) |

*Survey 1-4: Data not collected

### Supplement 3, Table 1. Adjusted prevalence ratio for VT carriage, analysis by survey within age group

| | Surv-1 | Survey-2<br>aPR[1]<br>95% CI; p-value | Survey-3<br>aPR<br>95% CI; p-value | Survey-4<br>aPR<br>95% CI; p-value | Survey-5<br>aPR<br>95% CI; p-value | Survey-6<br>aPR<br>95% CI; p-value | Survey-7<br>aPR<br>95% CI; p-value |
|---|---|---|---|---|---|---|---|
| **Infants 4-8wks (PCV unvaccinated)[2]** | - | - | - | - | Ref | 0.931<br>0.824-1.051; 0.252 | 0.980<br>0.908-1.058; 0.607 |
| **Children 18wks - 1yr (PCV vaccinated) [3]** | - | - | - | Ref | 0.998<br>0.924-1.078;<br>0.961 | 0.990<br>0.954-1.027; 0.586 | 0.986<br>0.957-1.016; 0.352 |
| **Children 2yrs (PCV vaccinated)[3]** | - | - | - | Ref | 0.939<br>0.866-1.018;<br>0.128 | 0.981<br>0.946-1.017; 0.295 | 0.979<br>0.950-1.008; 0.158 |
| **Children 3-7yrs (PCV vaccinated)[4]** | Ref | 1.008<br>0.959-1.059; 0.753 | 1.006<br>0.982-1.030; 0.631 | 0.994<br>0.977-1.012; 0.533 | 1.000<br>0.987-1.014;<br>0.955 | 0·991<br>0·980-1·003;<br>0·128 | 0·994<br>0·985- 1·004;<br>0·272 |
| **Children 3-10yrs (PCV unvaccinated) [4]** | Ref | 0·983<br>0·934-1·033; 0·491 | 0·980<br>0·953-1·006; 0·135 | 0·974<br>0·953-0·995;<br>0·016 | 0·966<br>0·941-0·991;<br>0·009 | 0·978<br>0·963-0·994;<br>0·006 | 0·991<br>0·979-1·002;<br>0·107 |
| **Adults 18-40yrs, HIV-infected (PCV unvaccinated) [4]** | Ref | 0·988<br>0·912-1·070; 0·763 | 0·991<br>0·956-1·026; 0·602 | 0·995<br>0·972-1·244;<br>1·019 | 0·983<br>0·964-1·002;<br>0·079 | 0·981<br>0·965-0·996;<br>0·016 | 0·983<br>0·969-0·998;<br>0·031 |

[1]aPR: adjusted prevalence ratio; adjusted for age (years)
[2]Recruitment of Infants 4-8 weeks started at survey 5
[3]Recruitment of children 18 weeks-2 years started at survey 4
[4]Recruitment for children 3-10 years and adults started at survey 1
